# Mouth magnetoencephalography: A unique perspective on the human hippocampus

**DOI:** 10.1101/2020.03.19.998641

**Authors:** Tim M. Tierney, Andrew Levy, Daniel N. Barry, Sofie S. Meyer, Yoshihito Shigihara, Matt Everatt, Stephanie Mellor, Jose David Lopez, Sven Bestmann, Niall Holmes, Gillian Roberts, Ryan M Hill, Elena Boto, James Leggett, Vishal Shah, Matthew J. Brookes, Richard Bowtell, Eleanor A. Maguire, Gareth R. Barnes

## Abstract

Traditional magnetoencephalographic (MEG) brain imaging scanners consist of a rigid sensor array surrounding the head; this means that they are maximally sensitive to superficial brain structures. New technology based on optical pumping means that we can now consider more flexible and creative sensor placement. Here we explored the magnetic fields generated by a model of the human hippocampus not only across scalp but also at the roof of the mouth. We found that simulated hippocampal sources gave rise to dipolar field patterns with one scalp surface field extremum at the temporal lobe and a corresponding maximum or minimum at the roof of the mouth. We then constructed a fitted dental mould to accommodate an Optically Pumped Magnetometer (OPM). We collected data using a previously validated hippocampal-dependent task to test the empirical utility of a mouth-based sensor, with an accompanying array of left and right temporal lobe OPMs. We found that the mouth sensor showed the greatest task-related theta power change. We also found that, as predicted by the simulations, the mouth sensor was anti-correlated with those on over the temporal lobes. We found that this sensor had a mild effect on the reconstructed power in the hippocampus (~10% change) but that coherence images between the mouth sensor and reconstructed source images showed a global maximum in the right hippocampus. We conclude that augmenting a scalp-based MEG array with sensors in the mouth shows unique promise for both basic scientists and clinicians interested in interrogating the hippocampus.

## 1. Introduction

Optically Pumped Magnetometers (OPMs) offer new ways to explore the magnetic fields generated by human brain function. Simulation (Boto *et al.*, 2016; Iivanainen *et al.*, 2017) and empirical recordings (Boto *et al.*, 2017) have shown that it is possible to realize a fivefold signal magnitude increase for cortical sources, simply because OPMs can be placed much closer to the head (with a separation between the sensors’ sensitive volume and the scalp of around 6mm) compared to their cryogenic counterparts (which require a separation of around 17-30mm). However, for the hippocampus and other sub-cortical structures, the relative change in distance (and hence performance gain) we expect with OPMs over cryogenic systems is smaller - a factor of 2 or less - than for neocortical sources. For this reason, the ability to further leverage the flexibility of OPM-placement to design arrays that are specifically sensitive to these deeper brain areas is desirable.

In this study we exploited the flexibility offered by OPMs to test whether there are other places, besides the scalp surface, one might usefully place sensors. We first examined, in simulation, the topographies of simulated magnetic fields due to hippocampal sources over both the scalp surface and the roof of the mouth. We found that a typical hippocampal generator gave rise to a scalp surface field extremum over the temporal lobe with a corresponding maximum or minimum at the roof of the mouth. We then built a sensor casing into a dental mould and explored the empirical utility of such an arrangement. Using a previously validated hippocampal-dependent task (Barry *et al.*, 2019*a*, 2019*b*), we assessed the change in theta power across sensors. We then tested the importance of this additional channel for source reconstruction. Finally, we used the temporal lobe array to construct a beamformer image and tested for regions of the brain which were coherent with the mouth channel.

## 2. Materials and methods

The study had two components, an initial simulation phase followed by the recording and analysis of empirical data.

### 2.1 Exploring fields due to hippocampal generators

We first used a single participant head-model to explore the field generated across the scalp and over the roof of the mouth by current sources on the hippocampal manifold. We used the individual cortical surface of the participant as extracted from Freesurfer (Dale *et al.*, 1999). This included two additional surfaces comprising the hippocampal envelopes (as described in (Meyer *et al.*, 2017)). The outer scalp and inner-skull meshes were based on the SPM inverse-normalized template meshes (Mattout *et al.*, 2007; Litvak *et al.*, 2011). We assumed the OPMs to be ideal point-source magnetometers with no orientation, position or gain errors. All lead-field calculations were based on the Nolte single-shell forward model (Nolte, 2003). To produce a scalp-level field map for each hippocampal source we computed point estimates that were oriented normal to the outer scalp surface and offset by 6.5mm from the surface in this direction. This resulted in 2562 samples of external (scalp) field for each source on the hippocampal envelope.

### 2.2 Empirical recordings

#### 2.2.1 Participants

One participant (male, aged 50 years) took part in the study. Data collection took place at the University of Nottingham, UK. The research protocol was approved by the University of Nottingham Medical School Research Ethics Committee and written informed consent was obtained from the participant. The data from the temporal channels of this subject formed part of the cohort of participants reported in (Barry *et al.*, 2019*b*).

#### 2.2.2 Mouth sensor holder

In order to record from the roof of the mouth, an intraoral appliance to hold the OPM sensor was constructed by S4S (UK) Limited (https://www.s4sdental.com/). Construction started with standard intraoral impressions of the upper and lower dental arches. The appliance (Figure 1) was constructed from 3mm Erkoloc-Pro (Erkodent Australia). This is a dual-laminate material composed of two individual thermoplastic layers that are chemically bonded: soft inner layer, helping to improve the comfort of the appliance, and a rigid outer layer that provides stiffness and is able to withstand forces from biting. The appliance was constructed on the upper dental arch which provided a stable base. The use of material with a soft compressible lining enabled us to comfortably engage the majority of the tooth surface while reducing movement and rotation, with minimal risk of the appliance being unremovable. The OPM was fully encapsulated by the appliance to minimise saliva contamination. The dual-laminate material was able to undergo the repeated disinfections needed to ensure hygiene without deterioration.

**Figure 1.**
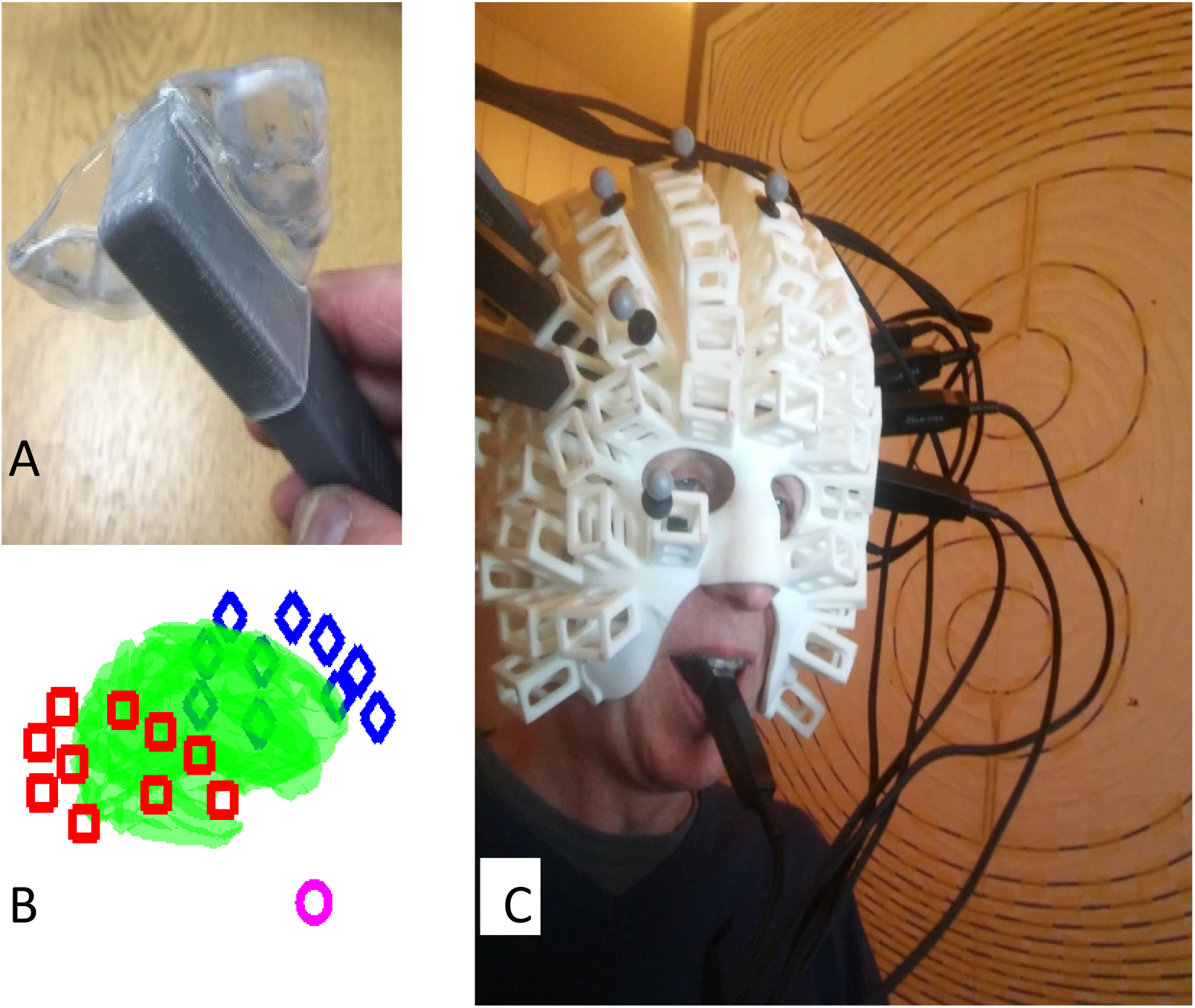
Experimental set-up. **A.** The custom translucent thermoplastic intraoral sensor holder to encapsulate the end of a Quspin Gen 1 sensor (grey). **B.** Distribution of the sensors with respect to the participant’s cortex (green). The mouth sensor is shown as a pink circle, right and left temporal lobe sensors are shown as red boxes and blue diamonds, respectively. **C.** The participant wearing a scanner-cast with the temporal lobe OPM array and the mouth sensor.

A limitation on how far into the mouth the appliance can be placed is imposed by the need to avoid activating the gag reflex by impinging on structures in the posterior portion of the oral cavity (soft palate, posterior of tongue, uvula, posterior wall of pharynx, palatoglossal and palatopharyngeal arches). We determined the posterior limit of the appliance to be just anterior of the soft palate. The border of the hard and soft palate is also clearly identifiable both intraorally and radiographically. Based on visual inspection (with approximately ±1cm of potential error) of the participant’s structural MRI brain scan, we estimated the location of the mouth sensor in native space. This corresponded to MNI coordinates x=−2.4, y=15, z=−103 and the orientation of its sensitive axis in MNI space to be described by the unit vector (0, 0.9885, −0.1513).

#### 2.2.3 Hippocampal-dependent task

We used a task known to be hippocampal-dependent, full details of which are described in (Barry *et al.*, 2019*a*, 2019*b*). In summary, the experimental task required the imagination of novel scenes in response to single-word cues, and there was an additional baseline condition involving counting. During scanning, experimental stimuli were delivered aurally via an MEG-compatible earbud using the Cogent toolbox (www.vislab.ucl.ac.uk/cogent.php), running in MATLAB. To prepare the participant for each trial type, they first heard either the word “scene” or “counting”. The participant immediately closed his eyes and waited for an auditory cue which was presented following a jittered delay of between 1300 and 1700 ms. During each scene trial, the participant had 3000ms to construct a novel, vivid scene in their imagination based on the cue (e.g. “jungle”). Each counting trial involved mentally counting in threes from a given number cue (e.g. “forty”) for 3000ms (beginning after the instruction had ended).

Data were recorded within a magnetically shielded room (MSR). The participant wore a 3D printed scanner-cast that accommodated 20 temporal lobe OPM sensors bilaterally and a single mouth OPM in its custom made holder. The participant’s head-movement was monitored using an Optitrack V120:Duo motion capture system (https://optitrack.com/) using an array of retro-reflective spheres attached to the scanner-cast (Figure 1C). OPM data were sampled at 1200 Hz using a 16-bit national instruments A/D converter. Data were recorded in 3 contiguous blocks and concatenated resulting in a total of 73 scene, and 68 counting, trials, each of 3000ms duration.

A bi-planar coil system (Holmes *et al.*, 2018) was used, in conjunction with a reference array (comprising 4 OPMs placed immediately behind the participant), to cancel any remaining static background field inside the MSR; specifically, reference array measurements enabled calculation of optimised coil currents to remove the static field, and its first order spatial derivatives over a central volume (Boto *et al.*, 2018; Tierney *et al.*, 2018).

All OPM data were first acausally filtered 1-8Hz using a 4^th^ order Butterworth filter. The data were then epoched into 3 second blocks based on digitally recorded triggers. The reference OPM array, and its temporal derivatives (i.e. 8 channels) were then used to regress any remaining environmental noise from the scalp OPM data on a trial by trial basis.

In order to verify that the data from the mouth sensor was qualitatively consistent with those from the temporal lobes we constructed time-frequency spectrograms of the difference between scene and counting trials at each sensor. We used the field-trip ((Oostenveld *et al.*, 2011), http://www.fieldtriptoolbox.org/) based multi-taper spectral estimate method (spm_eeg_specest_ft_mtmconvol.m) over 1-8Hz and 0-3000ms. We then used a paired sample t-test to compare between time-frequency bins in the two conditions of interest.

Based on our previous cryogenic MEG experiment using the same stimuli (Barry *et al.*, 2019*a*) we had a single hippocampal-specific time frequency window of interest of 0-3000ms and 4-8Hz (theta power). Based on the simulations predicting opposite hippocampal field extrema at mouth and temporal lobes, we first looked at the sensor level correlation (across all trials) of the mouth sensor and all temporal lobe sensors in this window. At the sensor level we then tested for the anticipated change in theta power between the 0-3000ms post-stimulus windows in counting and scene conditions. Data from each trial and channel were Hanning-windowed and band-pass filtered from 4-8Hz. We used a paired-sample t-test to look for power change between scene trials and counting trials (68 of each in order to equalise the comparison).

All subsequent processing was carried out in SPM (https://www.fil.ion.ucl.ac.uk/spm/) or DAISS (https://www.fil.ion.ucl.ac.uk/spm/ext/#DAiSS). We performed the same contrast (scene versus counting, 4-8Hz, 0-3000ms) at the source level (grid spacing 5mm) using an LCMV beamformer with automated Minka truncation (Minka, 2000) to produce volumetric whole-brain images. We used the multivariate implementation of the LCMV beamformer in DAISS to perform this univariate test. This returns a classical F statistic (which we report here) in the univariate case. In order to look for MEG sensors making the greatest impact at the hippocampus, we systematically removed one MEG sensor at a time from the analysis and calculated the mean F statistic within this structure. Channels that have a positive impact on the source reconstruction should produce a lower F-statistic in the hippocampus when removed from the source reconstruction.

Finally, we used a Dynamic Imaging of Coherent sources (DICs) beamformer with the mouth sensor (excluded from the source reconstruction) as the reference signal in order to create mouth-brain coherence images during the scene imagination condition. Covariance and coherence windows were 0-3000ms post cue onset, bandwidth was 4-8Hz and the grid spacing was 5mm. The resulting images were then smoothed to 15mm. In order to establish a significance threshold, we shuffled the mouth sensor trials and produced 100 (smoothed) coherence null images. Taking the maximum from each image established a null distribution which resulted in a coherence threshold corresponding to p<0.01 (whole-volume corrected).

## 3. Results

### 3.1 Fields due to hippocampal generators

We first explored the sensitivity of all possible extra-cranial recording positions to sources on the hippocampal envelope. The SPM-extracted scalp mesh covered the external scalp contours and was a closed-form, approximately elliptical, structure. The mesh passed below the occiput, travelled through the base of the spine and, following the roof of the mouth, emerged onto the scalp surface once again at the approximate level of the nasion (Figure 2A). At the source level, we used an individually segmented hippocampal surface for the single participant, with sources oriented normal to the hippocampal envelope (as in (Meyer *et al.*, 2017)). Based on sequentially positioning dipolar sources along this hippocampal model, we calculated the field magnitude at points on a shell displaced 6.5mm from (and normal to) the scalp surface as an estimate of measurable OPM signal (Figure 2B). The schematic in the centre of Figure 2 displays the viewpoint for the subsequent shells which show the external field due to hippocampal sources; note that the base of the shell approximately corresponds to the roof of the mouth.

**Figure 2.**
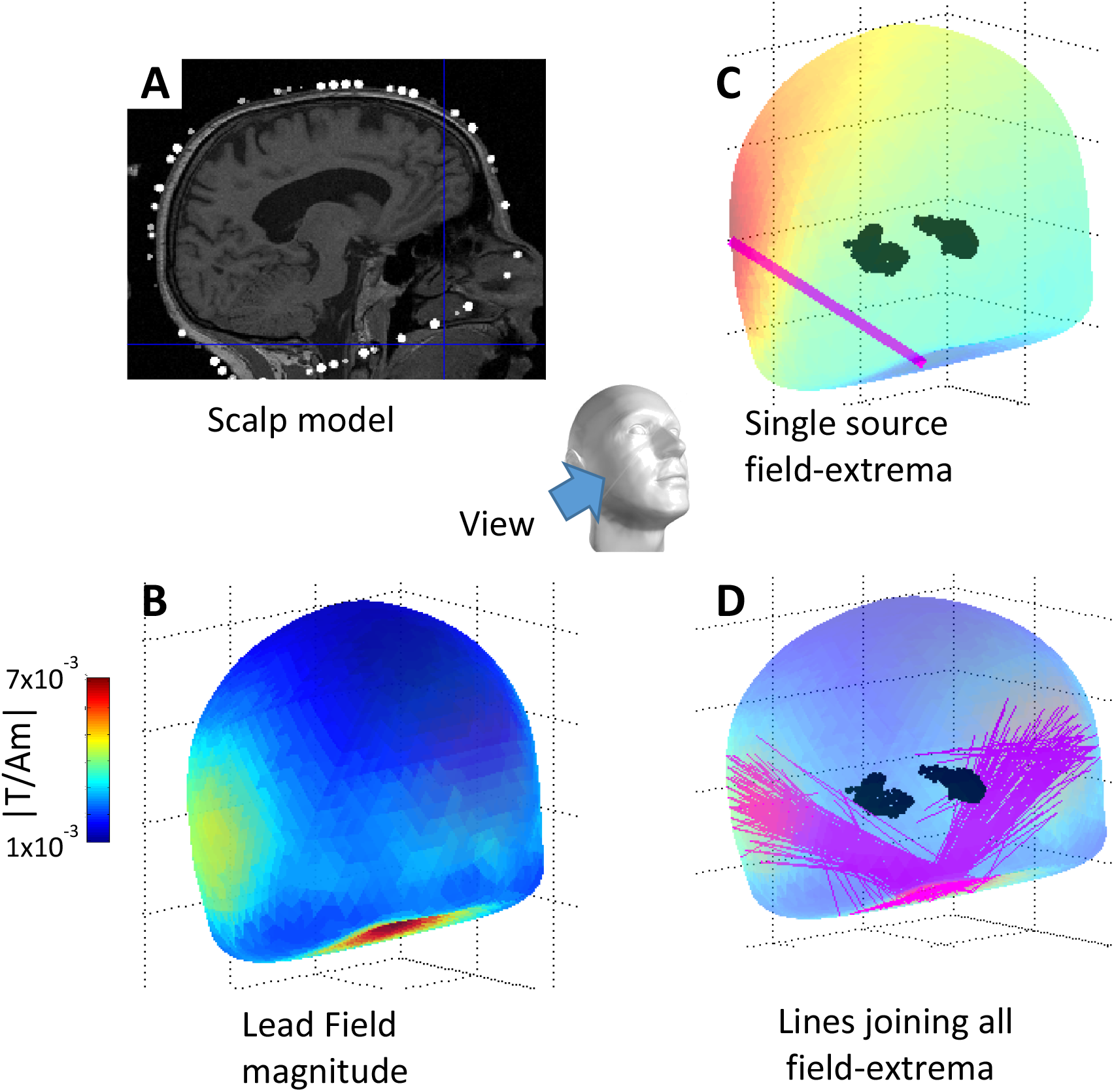
Exploring the lead-field pattern due to hippocampal sources. **A.** Sagittal section from the MRI brain scan of the participant showing the SPM-extracted scalp mesh (white dots) and its path along the roof of the mouth. The location of mouth sensor is shown by blue cross-hairs. **B.** The average field magnitude due to hippocampi on a shell displaced 6.5mm from the scalp surface (angle of view for B,C,D is shown in the central panel). Note the extrema at the temporal lobes and the roof of the mouth. **C.** The field due to a single right hippocampal source. The pink line joins the field extrema at the right temporal lobe and the roof of mouth **D.** The lines joining all field extrema for all hippocampal current elements. Note the clear pattern, with each hippocampal source giving rise to maximal (and opposing) field changes on one temporal lobe and the roof of the mouth.

It is clear from Figure 2B that the hippocampal generators gave rise to a large field magnitude on the temporal lobes, but an even larger field was expected at the roof of the mouth. It is instructive to examine the lines joining positive and negative field extrema due to each hippocampal dipolar source. Figure 2C displays the field due to a source on the right hippocampal envelope which showed a positive extremum on the right temporal lobe and a negative extremum at the roof of the mouth. These extrema were joined with a line. We then moved through each hippocampal source in turn and drew a line connecting the extra-cranial lead-field maxima and minima (Figure 2D). It is striking that the hippocampi generated fields that had extrema on the left and right temporal lobes (for left and right hippocampi respectively) with additional (anti-correlated) companion extrema at the roof of the mouth.

### 3.2 Empirical recordings

Based on the simulations described above, we proceeded to test the feasibility of taking measurements from within the mouth cavity while the participant performed the hippocampal-dependent scene imagination task (Barry *et al.*, 2019*a*,*b*).

Figure 3 shows the sensor level data. Panels A-C show that the mouth sensor recordings are qualitatively similar to the temporal lobes channels suggesting that we have access to neuronal (rather than tongue or other artefactual) recordings. In panel D the mean cross-correlation (4-8Hz, 0-3sec) between the time-series data from mouth sensor and left and right temporal sensors is shown. Note that as predicted (Figure 2) all temporal channels (but one) are negatively correlated with the mouth sensor in the (hippocampal related) 4-8Hz band.

**Figure 3.**
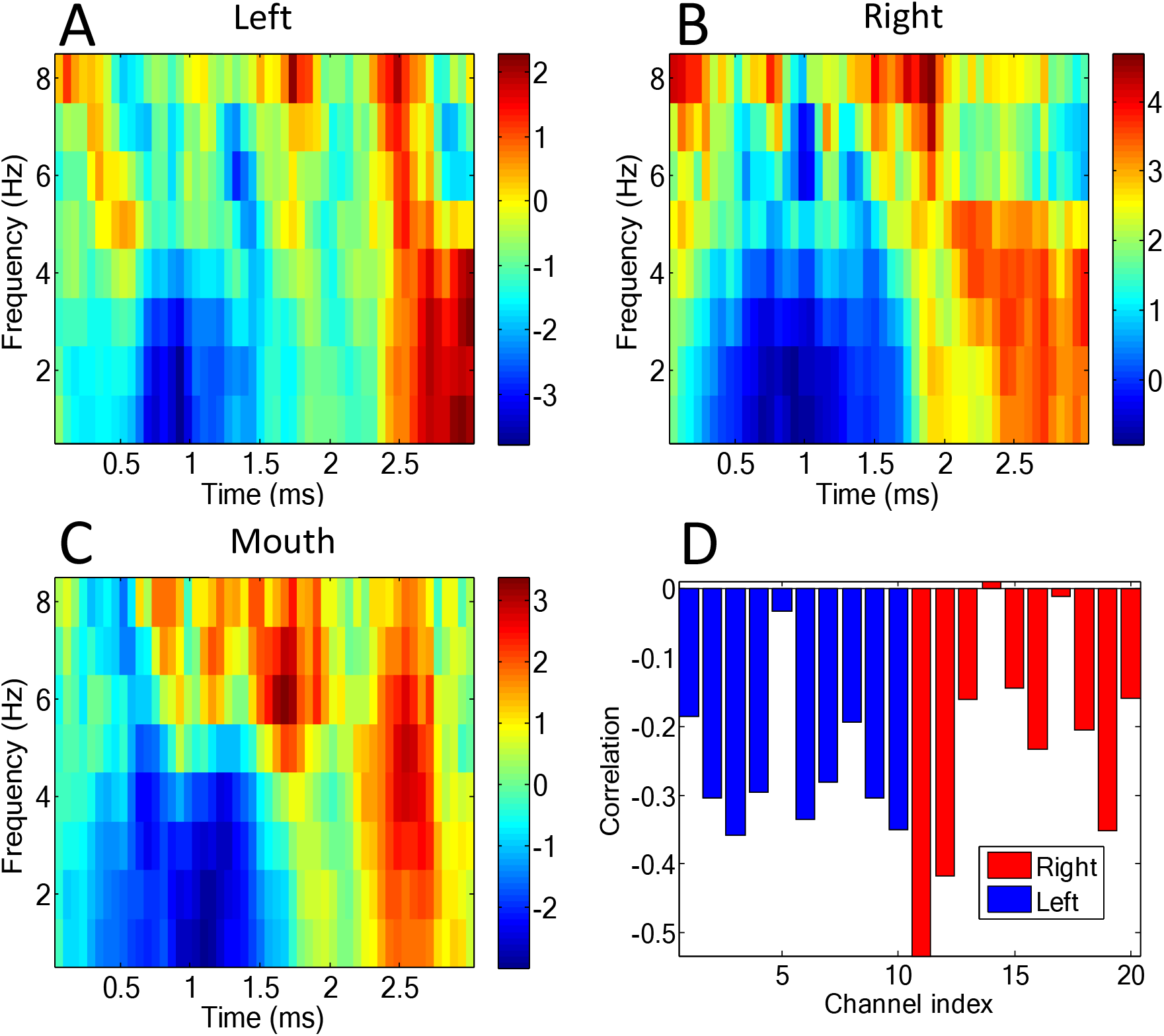
Initial sensor level validation. Panels **A** and **B** show time-frequency spectrograms (1-8Hz, 0-3sec) of the t-statistical difference between scene and counting conditions for representative left, right temporal channels. Panel **C** shows the same contrast at the mouth sensor. Panel **D** shows the average correlation coefficient between the time-series data (in the 4-8Hz band) from the mouth sensor and all left (blue) and right (red) temporal lobe channels.

Figure 4A shows the channel-level t-statistical power changes between scene imagination and counting conditions based on our prior hypothesis (4-8Hz, 0-3000ms (Barry *et al.*, 2019*a*)). Note that the largest absolute t-statistic (t=3.08, df=134, p<0.0025) occurred at the mouth sensor. The fact that this sensor, out of 21 sensors in total, showed the largest change is unlikely to have occurred by chance (p<0.0476). This suggests that, not only is the mouth sensor picking up useful signal, but this signal is strongly modulated by a stimulus we know engages the hippocampus.

**Figure 4.**
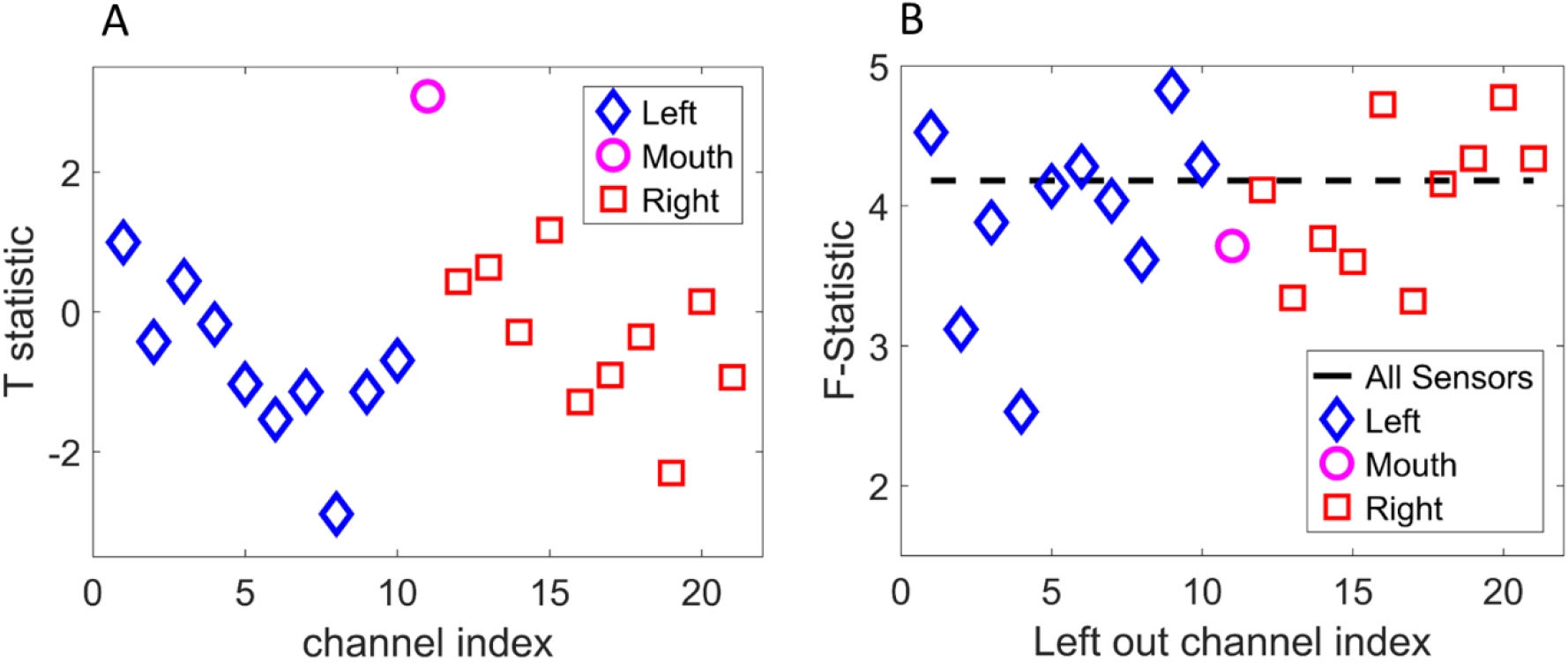
Channel-specific tests at sensor and source level. The mouth sensor, left, and right temporal lobe channels are depicted as a pink circle, blue triangles and red squares respectively. **A.** Single sensor-level two-sample tests on the theta power difference between scene imagination and counting trials. The largest task modulation (largest absolute t-statistic) is at the mouth sensor. **B.** F-statistic (relative power change) within the hippocampi when each measurement channel is excluded. The dotted line (baseline) indicates the F-statistic (power change) when using all channels. Removal of channels critical to the analysis should lead to a drop in power. Here we find that although the mouth sensor is important it is not as essential as some of the temporal lobe channels.

We then constructed a beamformer image of the contrast between scene imagination and counting (again over a 3 second window in the 4-8Hz band). In order to identify channels key to explaining experimental variance within the hippocampus, we re-ran the beamformer reconstruction, but each time omitted one of the measurement sensors. Channels which were key to explaining experimental variance should give rise to lower F-statistic when omitted. Figure 4B shows that the sensor that had the greatest impact on the amount of experimental variance explained was a channel on the left temporal lobe (channel 4). The impact of the mouth sensor on this analysis was modest (~10%). The fact that we observed maximal experimental modulation at the mouth sensor, but that it made a small contribution to the source imaging, suggested to us that the lead-fields for that sensor might be in error.

To further probe whether or not the signal from the mouth sensor was coming from the hippocampus we used DICs sources. This allowed us to identify which brain regions were most coherent with the mouth sensor. The advantage of this analysis is that it does not require an explicit sensitivity profile (or lead field), for the mouth sensor.

Figure 5 shows the DICs image of coherence between the mouth sensor and the beamformer source locations throughout the brain. We found the greatest coherence between the mouth sensor and source time-series within the beamformer image to be located in the right hippocampus (this was the global image maximum). Only one other peak survived the whole volume statistical correction (p<0.01) and this bordered primary motor and Brodmann Area 6.

**Figure 5.**
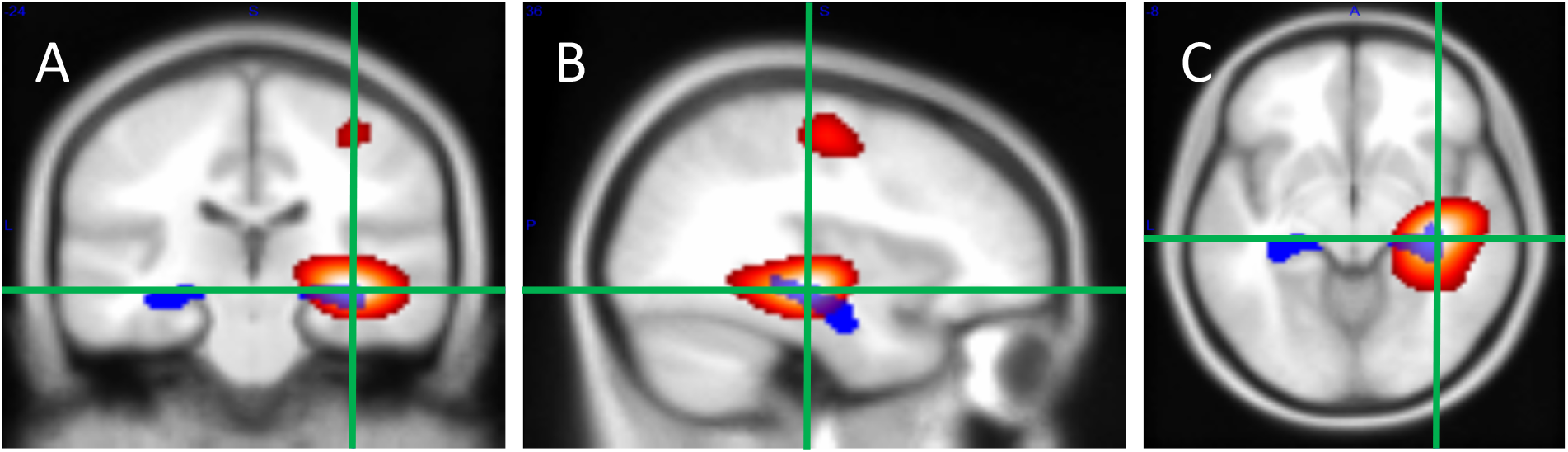
Coherence images from the OPM array whilst performing a scene imagination task superimposed on a template brain. Regions with high coherence to the mouth sensor are shown in hot colours (display threshold at p<0.01 (coherence=0.0969), whole volume corrected). Coronal **A**, sagittal **B** and axial **C** slices are shown at the global coherence peak (green cross-hairs) (coherence=0.1527, x=36.00 y=−24.00 z=−8.00). The AAL anatomical location of the hippocampi are shown in blue. Only two peaks are significant, the largest in the right hippocampus (on which the images are centred). The secondary peak (36.00 −16.00 54.0) is at the border of primary motor cortex and BA6.

## 4. Discussion

We showed in simulation that the inclusion of a mouth sensor could add a new dimension to on-scalp MEG measurements. The simulation predicted an enhanced sensitivity to hippocampal generators within the mouth and that the mouth signal should be anti-correlated with that from either temporal lobe. We then demonstrated this anti-correlation empirically (Figure 3D) and provided the first demonstration of a mouth sensor’s hippocampal selectivity, both spectrally (Figure 4A) and spatially (Figure 5), to the human hippocampus.

We were initially surprised by the insights from the simulation study which clearly identified the roof of the mouth as the site of magnetic field extrema due to sources at the hippocampal surface. There are clear parallels here with the use of sphenoidal electrodes in EEG (Jones, 1951; Pampiglione & Kerridge, 1956) to access the base of the brain. Our simulation also suggested that each hippocampus should produce a unilateral temporal lobe extremum in conjunction with that found in the mouth. Reassuringly, recent simultaneous intracerebral electrophysiological and MEG recordings (Pizzo *et al.*, 2019) have led to similar observations, with the invasively recorded hippocampal source giving rise to a strong, yet unilateral, temporal lobe signal.

The hippocampus is the target of the majority of adult epilepsy surgeries (Margerison & Corsellis, 1966) (Walker, 2015) and is heavily implicated in the progression of several forms of dementia (Huijbers *et al.*, 2015; Buzsáki, 2015). This vulnerable brain structure is, therefore, an important focus for any non-invasive clinical imaging system. However, the main sensitivity benefit of OPMs over SQUID MEG systems is cortically focussed, with idealized sensitivity gains falling from five-fold cortically to two-fold for deeper structures (Boto *et al.*, 2016; Iivanainen *et al.*, 2017). Our findings show that mouth-based sensor arrays for MEG could potentially further enhance sensitivity to deep structures like the hippocampus.

Clinically, the ability to estimate electrical activity from the hippocampus non-invasively using mouth-based arrays would pose a much reduced risk compared to the surgical implantation of electrodes within the hippocampus, which is currently best-practice in cases when the source of the seizure focus is uncertain. Eliminating the need for this additional operation could significantly shorten the pathway to surgery to remove the aberrant seizure-inducing tissue. This study made employed Gen 1 Quspin sensors and we only made use of measurements from one axis (axial to the sensor body). At present, the set-up needed to use a intraoral OPM sensor is cumbersome and uncomfortable; however, OPMs are continuing to decrease in size (Alem *et al.*, 2014; Osborne *et al.*, 2018). We hope that with improved sensor technology (and possibly by measuring a field from two orthogonal directions simultaneously) small mouth-based arrays might be possible in future.

Here we found that although the mouth sensor explained the most experimental variance (Figure 4A) it was not the most important sensor for the source level analysis (Figure 4B). A possible explanation for this is that the lead-fields for the mouth sensor may have been sub-optimal. The sensor position and orientation were estimated by visual inspection and could be in error by around 1 cm. Furthermore, it was not possible to orient the sensitive axis of the sensor in the mouth at the ideal angle used in the simulations (approximately 30 degrees offset). In future studies, this could be improved by utilizing the additional measurement orientation offered by many OPMs (axial as well as tangential to the sensor body) to obtain better sampling of brain signals recorded using intraoral sensors.

In conclusion, we have demonstrated the feasibility of acquiring meaningful data using a scalp-array of OPM sensors augmented by an intraoral sensor. This intraoral sensor provides higher signal to noise than the temporal lobe sensors and is most coherent with the signal in the hippocampus. These results illustrate the potential that this approach holds for interrogating deep structures like the hippocampus in basic science and clinical studies.

## Acknowledgements

This work was supported by a Wellcome collaborative award to G.R.B., M.J.B. and R.B. (203257/Z/16/Z, 203257/B/16/Z), and a Wellcome Principal Research Fellowship to E.A.M. (210567/Z/18/Z). The Wellcome Centre for Human Neuroimaging is supported by a Centre Award from Wellcome (203147/Z/16/Z). We would like to thank S4S for generously providing the sensor housing and their time free of charge. The scanner-casts were designed and built by Mark Lim at Chalk Studios Ltd.

## References

Alem O, Benison AM, Barth DS, Kitching J & Knappe S (2014). Magnetoencephalography of Epilepsy with a Microfabricated Atomic Magnetrode. J Neurosci 34, 14324–14327.

Barry DN, Barnes GR, Clark IA & Maguire EA (2019a). The neural dynamics of novel scene imagery. J Neurosci2497–18.

Barry DN, Tierney TM, Holmes N, Boto E, Roberts G, Leggett J, Bowtell R, Brookes MJ, Barnes GR & Maguire EA (2019b). Imaging the human hippocampus with optically-pumped magnetoencephalography. Neuroimage 116192.

Boto E, Bowtell R, Krüger P, Fromhold TM, Morris PG, Meyer SS, Barnes GR & Brookes MJ (2016). On the Potential of a New Generation of Magnetometers for MEG: A Beamformer Simulation Study. ed. Johnson B. PLoS One 11, e0157655.

Boto E, Holmes N, Leggett J, Roberts G, Shah V, Meyer SS, Muñoz LD, Mullinger KJ, Tierney TM, Bestmann S, Barnes GR, Bowtell R & Brookes MJ (2018). Moving magnetoencephalography towards real-world applications with a wearable system. Nature; DOI: 10.1038/nature26147.

Boto E, Meyer SS, Shah V, Alem O, Knappe S, Kruger P, Fromhold TM, Lim M, Glover PM, Morris PG, Bowtell R, Barnes GR & Brookes MJ (2017). A new generation of magnetoencephalography: Room temperature measurements using optically-pumped magnetometers. Neuroimage 149, 404–414.

Buzsáki G (2015). Hippocampal sharp wave-ripple: A cognitive biomarker for episodic memory and planning. Hippocampus 25, 1073–1188.

Dale AM, Fischl B & Sereno MI (1999). Cortical Surface-Based Analysis. Neuroimage 9, 179–194.

Holmes N, Leggett J, Boto E, Roberts G, Hill RM, Tierney TM, Shah V, Barnes GR, Brookes MJ & Bowtell R (2018). A bi-planar coil system for nulling background magnetic fields in scalp mounted magnetoencephalography. Neuroimage 181, 760–774.

Huijbers W, Mormino EC, Schultz AP, Wigman S, Ward AM, Larvie M, Amariglio RE, Marshall GA, Rentz DM, Johnson KA & Sperling RA (2015). Amyloid-β deposition in mild cognitive impairment is associated with increased hippocampal activity, atrophy and clinical progression. Brain 138, 1023–1035.

Iivanainen J, Stenroos M & Parkkonen L (2017). Measuring MEG closer to the brain: Performance of on-scalp sensor arrays. Neuroimage 147, 542–553.

Jones DP (1951). The EEG society (The Electroencephalographic Society). September 30th. 1950 The National Hospital, Queen Square, W.C.I., London. Electroencephalogr Clin Neurophysiol 3, 100–101.

Litvak V, Mattout J, Kiebel S, Phillips C, Henson R, Kilner J, Barnes G, Oostenveld R, Daunizeau J, Flandin G, Penny W & Friston K (2011). EEG and MEG data analysis in SPM8. Comput Intell Neurosci; DOI: 10.1155/2011/852961.

Margerison JH & Corsellis JA (1966). Epilepsy and the temporal lobes. A clinical, electroencephalographic and neuropathological study of the brain in epilepsy, with particular reference to the temporal lobes. Brain 89, 499–530.

Mattout J, Henson RN & Friston KJ (2007). Canonical Source Reconstruction for MEG. Comput Intell Neurosci 2007, 1–10.

Meyer SS, Rossiter H, Brookes MJ, Woolrich MW, Bestmann S & Barnes GR (2017). Using generative models to make probabilistic statements about hippocampal engagement in MEG. Neuroimage 149, 468–482.

Minka TP (2000). Automatic choice of dimensionality f o r PCA. Available at: https://vismod.media.mit.edu/tech-reports/TR-514.pdf [Accessed April 29, 2019].

Nolte G (2003). The magnetic lead field theorem in the quasi-static approximation and its use for magnetoenchephalography forward calculation in realistic volume conductors. Phys Med Biol 48, 3637–3652.

Oostenveld R, Fries P, Maris E & Schoffelen J-M (2011). FieldTrip: Open Source Software for Advanced Analysis of MEG, EEG, and Invasive Electrophysiological Data. Comput Intell Neurosci 2011, 1–9.

Osborne J, Orton J, Alem O & Shah V (2018). Fully integrated, standalone zero field optically pumped magnetometer for biomagnetism. In Steep Dispersion Engineering and Opto-Atomic Precision Metrology XI, p. 51. SPIE. Available at: https://www.spiedigitallibrary.org/conference-proceedings-of-spie/10548/2299197/Fully-integrated-standalone-zero-field-optically-pumped-magnetometer-for-biomagnetism/10.1117/12.2299197.full [Accessed April 5, 2019].

Pampiglione G & Kerridge J (1956). ABNORMALITIES FROM THE TEMPORAL LOBE STUDIED WITH SPHENOIDAL ELECTRODES. J Neurol Neurosurg Psychiat; DOI: 10.1136/jnnp.19.2.117.

Pizzo F, Roehri N, Medina Villalon S, Trébuchon A, Chen S, Lagarde S, Carron R, Gavaret M, Giusiano B, McGonigal A, Bartolomei F, Badier JM & Bénar CG (2019). Deep brain activities can be detected with magnetoencephalography. Nat Commun 10, 971.

Tierney TM, Holmes N, Meyer SS, Boto E, Roberts G, Leggett J, Buck S, Duque-Muñoz L, Litvak V, Bestmann S, Baldeweg T, Bowtell R, Brookes MJ & Barnes GR (2018). Cognitive neuroscience using wearable magnetometer arrays: Non-invasive assessment of language function. Neuroimage 181, 513–520.

Walker M (2015). Hippocampal Sclerosis: Causes and Prevention. Semin Neurol 35, 193–200.

